# Generation of broad-spectrum vaccine platforms against tick-borne flaviviruses using ancestral sequence reconstruction

**DOI:** 10.1101/2025.10.31.685878

**Authors:** Chasity E. Trammell, Brooke M. Enney, Michelle J. Savran, Brian J. Geiss, Gregory D. Ebel

## Abstract

Tick-Borne Flaviviruses (TBFs) pose a significant global health threat, but few vaccines exist against these viral pathogens. Currently approved vaccines against TBFs are only protective against specific virus species, leaving the population at risk for infection by many existing and emerging TBFs. The development of vaccines that provide broad protection against a range of related TBFs has the potential of reducing disease burden against known and unknown viral pathogens. In this study we utilized ancestral sequence reconstruction (ASR) to design recombinant antigens of extinct ancestral viruses to serve as vaccines against multiple modern day TBFs. We reconstructed a common ancestral envelope (E) antigen that possesses high sequence and structural identity to several modern-day TBF pathogens and engineered this ancestral E antigen sequence into an attenuated Sindbis virus vaccine platform. We observed that this Sindbis virus was replication competent and generated sustained antigen expression *in vitro*. Immunizing mice with this vaccine candidate resulted in the production of significant neutralizing antibody active against virulent Powassan (POWV) and Langat (LGTV) viruses. Antibody neutralization correlated well with decreased clinical signs and increased survival from virulent POWV challenge. These data indicate that a common ancestral antigen from TBFs can provide protection against multiple modern day TBFs, and that the ASR approach may provide a valuable new pathway to safe and effective pan-TBF vaccines.

## INTRODUCTION

Tick-borne flaviviruses (TBFs) encompass a range of viruses that can cause severe encephalitic (e.g. Powassan virus [POWV], tick-borne encephalitic virus [TBEV], Louping ill virus [LIV]) and hemorrhagic (e.g. Alkhurma hemorrhagic fever virus [AHFV], Omsk hemorrhagic fever virus [OHFV], Kyasanur Forest disease virus [KFDV]) diseases. The risk of exposure to many of these viruses has increased over the past decade [1–3]. This increased hazard risk is due to environmental and human factors that promote exposure to vector-competent ticks among historically seronegative human populations [4–7]. These factors, which include dynamic environmental temperature and seasonal weather patterns, changes in land use, and altered distribution and density of vertebrate host populations, have promoted expansions of tick habitation range and feeding behavior to enhance virus transmission [8]. While the incidence and geographic distribution of TBFs have expanded, advancements in preventative and prophylactic treatment options have not, and thus remain limited.

TBEV is the only pathogen within the TBF clade with approved vaccines that can prevent or limit severe disease [9]. There are no clinically available vaccines for other TBFs (POWV, LIV, AHFV, OHFV or KFDV), leaving significant gaps in protection for large portions of the at-risk human population. Notably, while available and successful in addressing TBEV in Eurasia, current vaccines possess complicated administration schedules and limited efficacy in preventing infection [10]. Because of the hinderances and limitations that accompany the current vaccine approaches for addressing TBFs, there is a pressing need for development of novel approaches for vaccine antigen design. In particular, given the level of cross-neutralization that is observed between TBFs in TBEV [11] or POWV [12], an approach that aims at generating broad-spectrum protection against entire viral clades would be highly beneficial.

The traditional model of “one-pathogen/one vaccine” has generated several outstanding vaccines that have substantially reduced disease burden and incidence worldwide; however, a hindrance of such approach is that the number of emerging viruses outpaces the development of new vaccines, increasing costs and reducing effective vaccination rates. Due to these challenges, there has been a push to move towards developing broadly efficacious vaccines to get away from the “one-pathogen/one vaccine” paradigm. One approach that has demonstrated feasibility is designing viral antigens using ancestral sequence reconstruction (ASR), a bioinformatics approach commonly used in paleogenomics that infers nucleic- or amino-acid sequences based on the phenotypic properties of known modern-day sequences [13, 14]. Briefly, ASR uses multiple sequence alignments (MSA) of related modern-day sequences to generate a phylogenetic tree where the resulting horizontal comparisons between the modern-day sequences are used to derive predicted ancestral sequences for nodes within the tree. Predicted residues in each node are selected based on traditional maximum likelihood or Bayesian methods that ultimately designate a most likely residue, and thus protein sequence, based on existing, modern-day sequences [14, 15]. The advantage of reconstructing ancestral sequences is that we can investigate extinct sequences that may have existed at one point, but are antigenically related to modern-day sequences. Current approaches that rely on consensus sequences provide a partial view of sequence evolution and diversity dependent solely on the sequence alignments of modern-day sequences and may not be immunologically cross-reactive.

ASR has been successful in predicting extinct protein sequences and rescue of functional proteins such as those associated with thermostability [16] and enzymatic function [17] within bacteria species. This has enabled the characterization of proteins that would have otherwise been lost to time. Because of its success in studying extinct proteins associated with host function, further investigation has evaluated its viability for protein design in other fields such as vaccine development. For example, ASR has been used for antigen design and vaccine development against H5N1 influenza viruses (18). Ducatez et. al. demonstrated that vaccination against the extinct HA and NA proteins via ASR resulted in high neutralizing titers and protection against lethal viral challenge [18]. Similarly, ancestral sequence analysis and reconstruction is one approached used in the study of HIV evolution [19] and development of new vaccine candidates [20]. It has not been demonstrated if ASR could be used to design vaccines a common ancestor of different modern-day viruses within a related virus clade like TBFs.

Accordingly, we used ASR to reconstruct the envelope protein of the TBF common ancestor as a candidate antigen for the development of a broad-spectrum vaccine. We successfully inserted our ancestral antigen into an alphavirus-based infectious clone to function as a self-replicating vaccine platform. In a C57BL/6J mouse model, we observed no adverse effects following vaccination, and significant neutralizing titers against Powassan (POWV) and Langat (LGTV) viruses were generated. These findings suggest the ancestral envelope antigen was able to induce broad-spectrum neutralizing antibodies that reduce virus infectivity. We demonstrated that higher neutralizing antibody titers were associated with decreased morbidity and mortality in mice during lethal POWV challenge. Overall, our data suggests that ASR may be a viable and effective approach for designing broad-acting vaccines that target whole clades of related viruses.

## RESULTS

### Phylogenetic analysis and reconstruction of TBF ancestral envelope sequences

To generate an ancestral envelope antigen sequence for TBF envelope protein, a phylogenetic tree was produced from envelope amino acid sequences of TBFs available (**Table S1**) using MEGA11 [21]. The putative ancestral sequences were calculated for all internal nodes and predicted for the root of phylogeny (node 19). West Nile virus (NY-99), a mosquito-borne flavivirus, was set as the outgroup to quantify divergence of TBFs from other members of the *Flaviviridae* genus (**Fig. 1A**). Node 19, the furthest common ancestor for the TBF clade, was selected as the antigen sequence for our vaccine studies. Sequences for all nodes within the TBF phylogenetic tree are provided in Table S1. We constructed a folded protein model of our node 19 sequence using AlphaFold2 and aligned the node 19 sequence with assorted modern day TBF envelope proteins (**Fig. 1B**) [22]. Node 19 (green) shared sequence and structural similarities with the known sequences of envelope proteins like POWV (red) (**Fig. 1C**). Our initial computational analysis resulted in the reconstruction of an amino acid sequence for the TBF envelope protein that will function as our target antigen.

**Figure 1:**
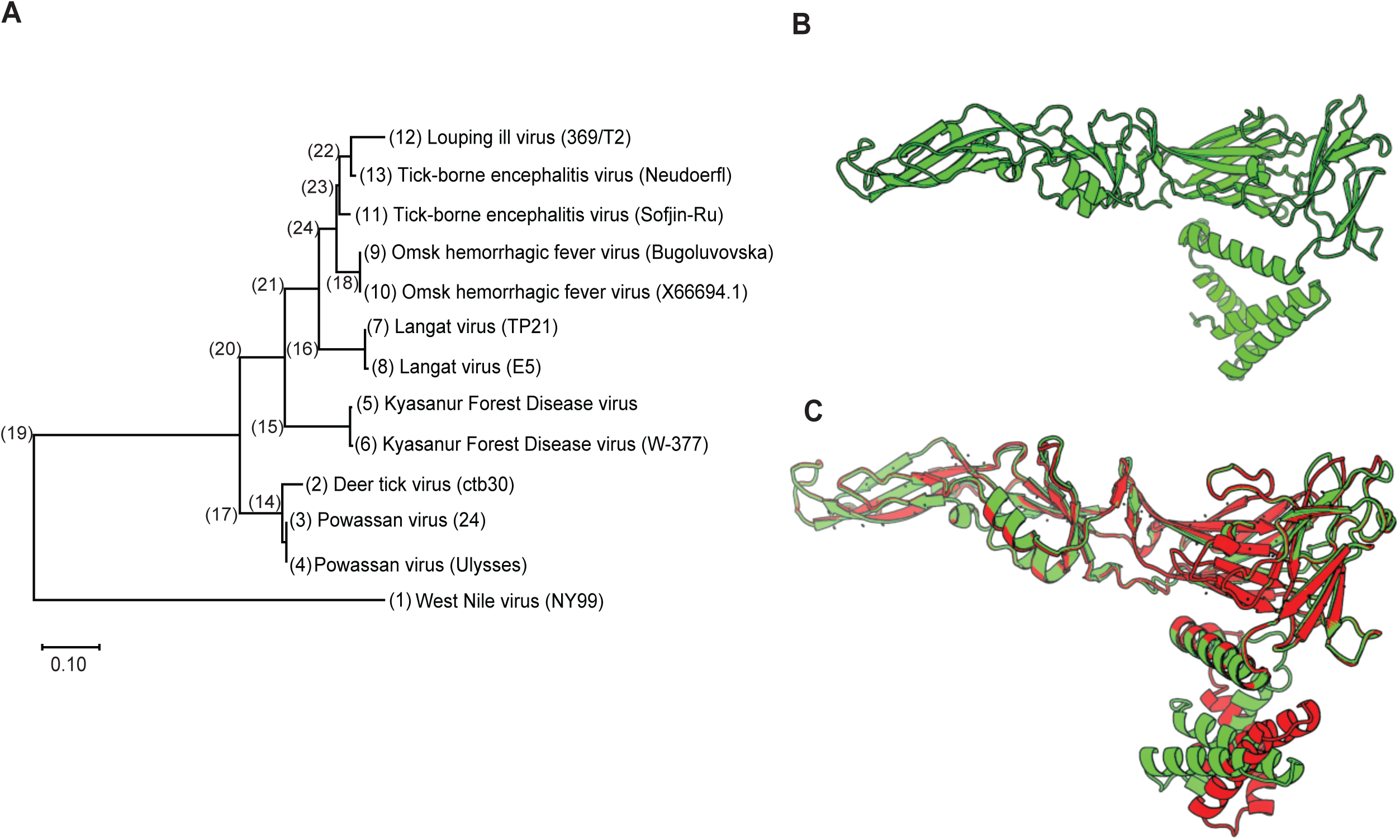
Phylogenetic relationship of modern-day tick-borne flaviviruses (TBFs) and inference of ancestral sequence of envelope protein for vaccine antigen design. (A) Sequences from multiple modern-day tick-borne flaviviruses were obtained from GenBank and phylogenetic relationship was inferred by using the Maximum Likelihood (ML) method and JTT matrix-based model (MEGA11). Phylogenetic tree analysis was rooted to West Nile virus (NY99), a mosquito-borne virus, as the designated outgroup for inference of internal node sequences. (Scale bar, 0.10 substitutions per site). Sequences used are provided in Table S1. (B) Computational protein folding model of the farthest common TBF ancestor (node 19) was generated using AlphaFold2. (C) Overlay of computationally-derived folded protein models for common TBF ancestor (green) with modern day POWV envelope protein (red).

### Construction of alphavirus expression platform of ancestral envelope protein as vaccine antigen

We used an established Sindbis virus (Family: *Togaviridae*) vector (pBG167) as our vaccine platform to take advantage of the viruses’ replication capacity and ease of manipulation [23]. We inserted the open reading frame for node 19 ancestral TBF E antigen (ASR TBF E) into a duplicated 3’ subgenomic promoter (3’ SGP) possessing a unique XbaI site in the pBG167 double subgenomic Sindbis virus expression plasmid via Gibson Assembly (**Fig. 2A**) [23]. To rescue antigen-expressing infectious virus, 125 ng of each plasmid construct (SINV WT and SINV +E) was transfected into BHK-21 cells using Lipofectamine 3000 in 6-well plates. Transfected cells were washed and media replaced with fresh growth media 4 hours post-transfection. Supernatant was harvested at approximately 4 days post transfection once >50% CPE was observed. This initial p0 stock was propagated in BHK-21 cells once (p1) to serve as our vaccine stocks.

**Figure 2:**
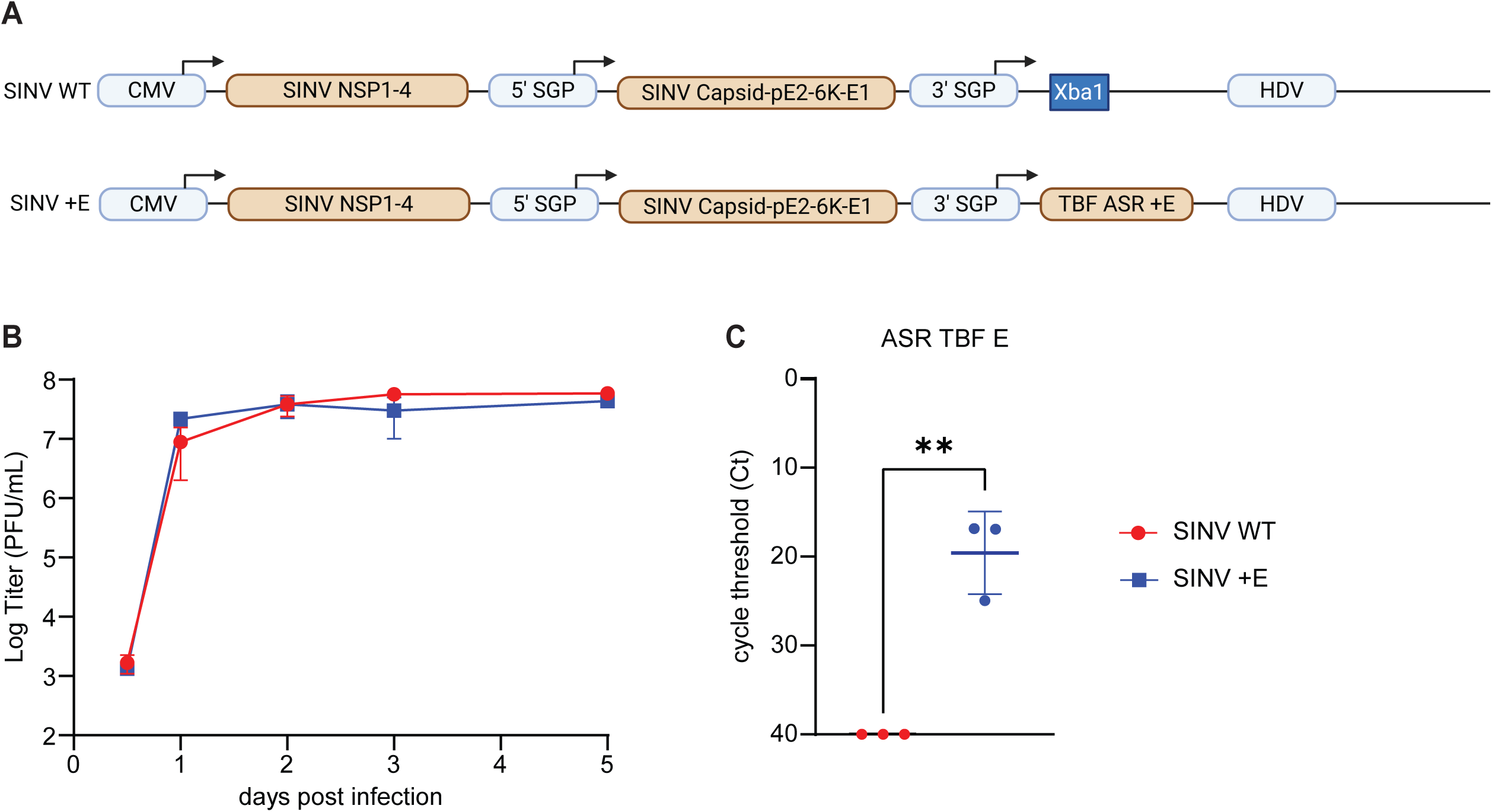
Plasmid construct and characterization of alphavirus-based vaccine platform. (A) Plasmid constructs were designed to function as vaccine platforms for high level expression of ancestral TBF envelope antigen using an alphavirus infectious clone platform (SINV WT). A gBlock consisting of the TBF +E sequence was inserted into a Sindbis virus infectious clone following a 3’ subgenomic promoter (3’ SGP) via restriction enzyme digestion and Gibson Assembly (SINV +E). (B) Viral titers of passage 1 (p1) from plasmid-transfected BHK-21 cells. BHK cells were infected at MOI=0.01. Supernatant was collected at indicated times and viral titers were determined by standard plaque assay. (C) Cycle threshold (Ct) values of p1 SINV WT (red) and SINV +E (blue) stocks probing for the ancestral TBF E (ASR TBF E).

We characterized the viral growth kinetics and antigen detection of our p1 vaccine stocks. BHK-21 cells were infected (MOI=0.01) and supernatant was harvested over 5 days post infection and quantified by standard plaque assay (**Fig. 2B**). We observed no difference in viral growth kinetics between backbone (SINV WT) and antigen-expressing viruses. Additionally, we confirmed stability and sufficient detection of our ancestral envelope gene by qRT-PCR (**Fig. 2C**) in our p1 vaccine stocks. Taken together, these results indicate that our insert is stable *in vitro* and can be sufficiently expressed using an alphavirus-based system.

### Vaccination against ancestral envelope antigen generates cross-reactive neutralizing antibodies

We assessed the safety and immunogenicity of the SINV +E vaccine candidate in a mouse model (**Fig. 3**). Briefly, female C57BL/6J mice were subcutaneously (SC) inoculated with 10^5^ PFU SINV +E or SINV WT (mock-vaccinated) diluted in 50 μL PBS or mock inoculated with 50 μL PBS. Mice were boosted with the same dose and route at 21 days-post-vaccination. Weight, clinical score, and survival were monitored over five weeks to assess vaccine safety. No variations in weight (**Fig. S1A**) nor clinical signs (**Fig. S1B**) were observed during the 35-day vaccine administration timeframe. The observed positive safety profile led us to pursue further characterization of the immunogenicity of our vaccine construct.

**Figure 3:**
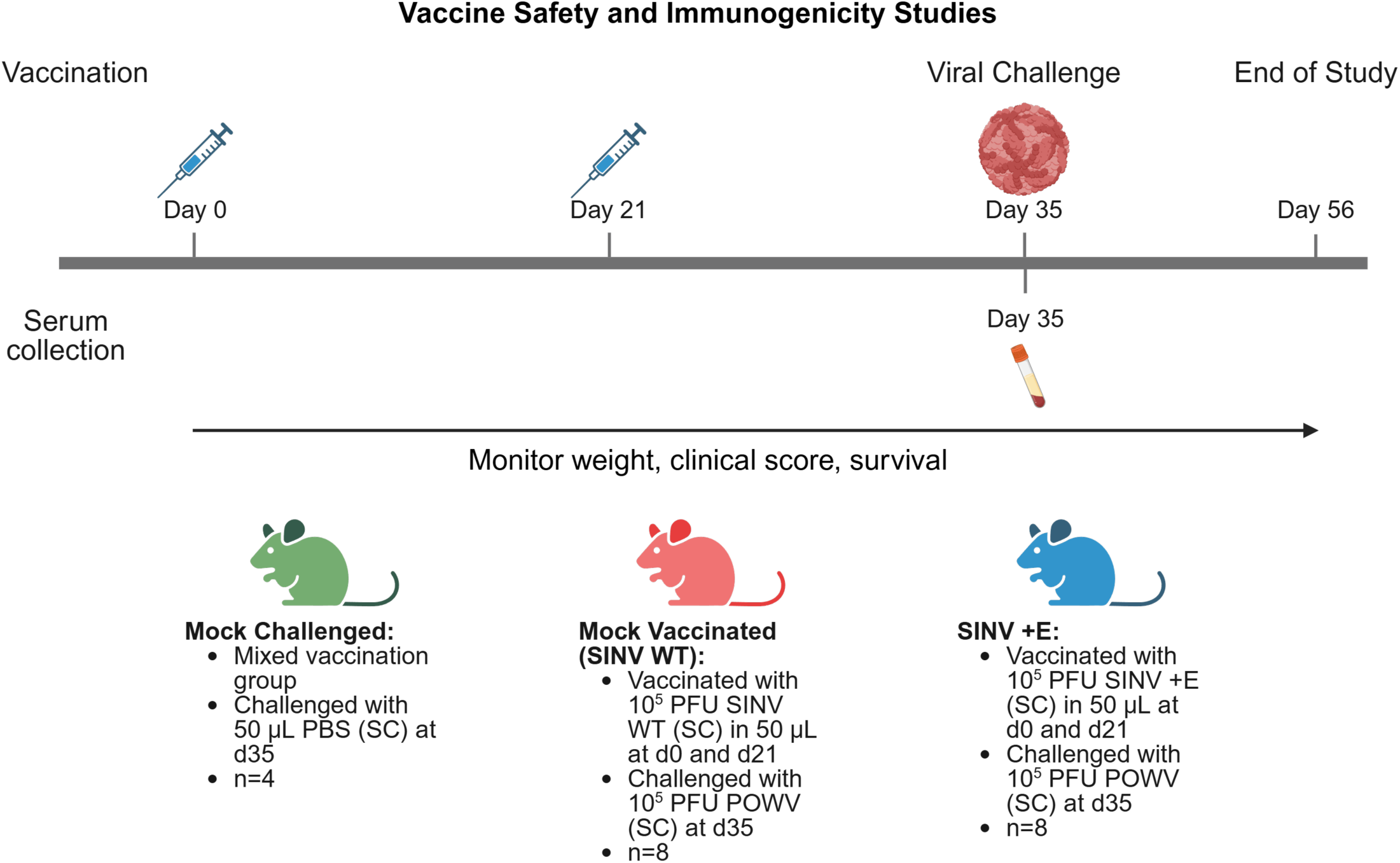
Assessment of vaccine safety and immunogenicity of SINV +E vaccine in C57BL/6J mice. Overview of animal studies in C57BL/6J mice. Female mice (n=8; 9 weeks old) were vaccinated with 10^5^ PFU of SINV +E (blue) subcutaneously (SC) in 50 μL solution at day 0 and day 21. Controls include mice vaccinated with 10^5^ PFU SINV WT (mock-vaccinated; red) and PBS (mock challenged; green). Whole blood was harvested and sera isolated by cardiac puncture at day 35 to assess neutralizing antibody titers. At day 35 mice are challenged with a lethal dose of 10^5^ PFU of Powassan virus SC diluted in 50 μL solution PBS. Weight, clinical score, and survival were monitored over 21 days post-challenge until end of study. Figure created in https://BioRender.com.

To assess whether SINV+E generated neutralizing antibodies that reduce viral infectivity of modern day TBFs, whole blood was collected via cardiac puncture 35 days post initial vaccination from a subset of vaccinated mice. Sera were isolated from whole blood samples, and neutralization was assessed against POWV, LGTV, and a chimeric LGTV infectious clone expressing the structural prM and E proteins of the BSL4 agent tick-borne encephalitis virus (TBEV) [24]. We also assessed neutralization against WNV as a control. We observed high neutralizing titers against POWV and LGTV (**Fig. 4A-B**) in SINV +E vaccinated mice compared to mock-vaccinated mice. We observed no limited neutralization against TBEV structural proteins (**Fig. 4C**) comparable to neutralizing titers against WNV (**Fig. 4D**). This suggests that while our vaccine candidate showed neutralizing capacity against evolutionarily divergent TBFs like POWV and LGTV, it was not able to generate antibody that could neutralize across the entire TBF clade.

**Figure 4:**
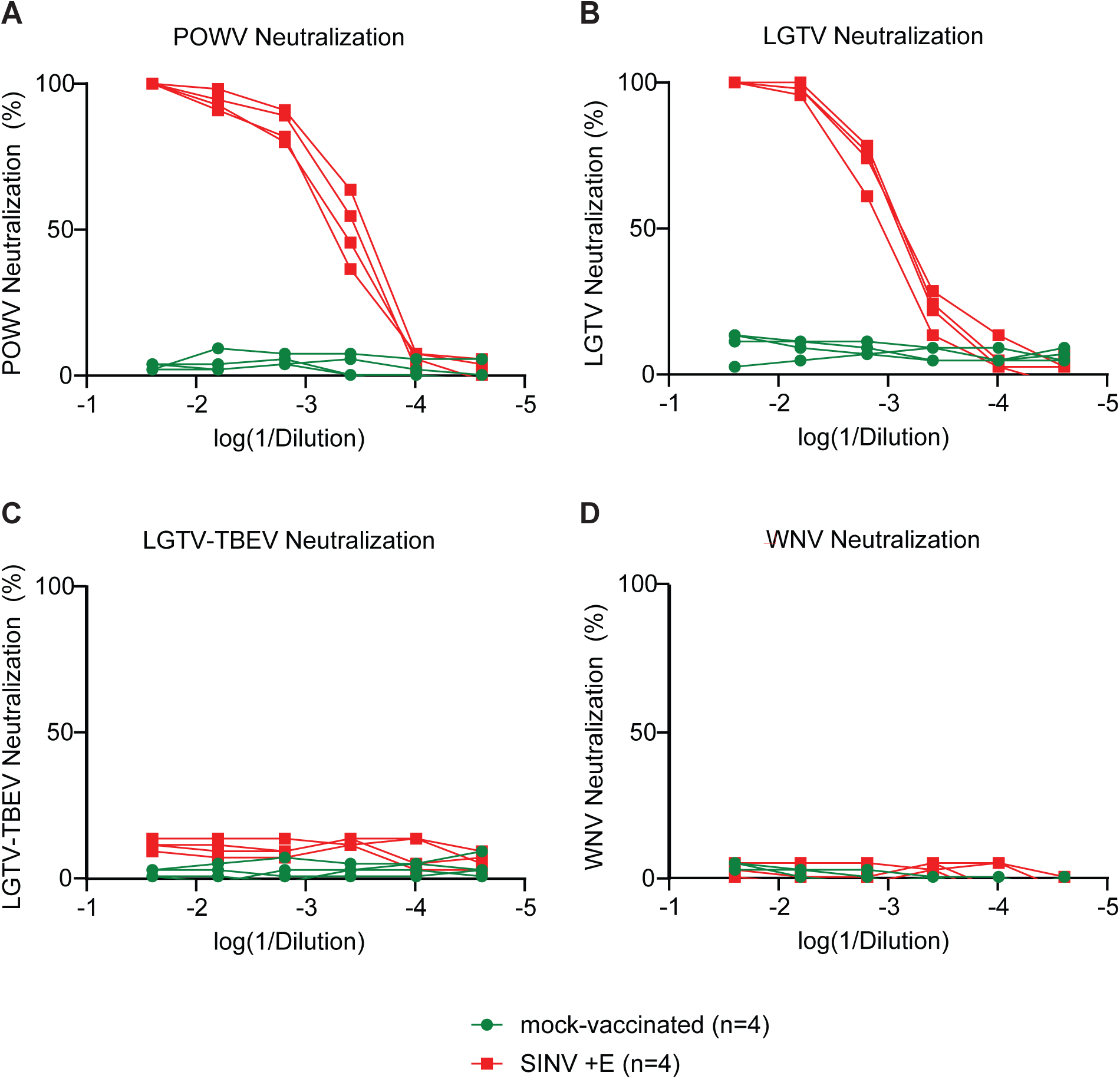
SINV +E vaccination generates neutralizing antibodies against Powassan and Langat viruses but not against TBEV. PRNTs performed on BHK-21 cells examined neutralization capacity of sera harvested at day 35 against (A) POWV, (B) LGTV, (C) LGTV-TBEV, and (D) WNV. Sera was collected from mock-vaccinated (green) and SINV +E vaccinated (red) mice. Each curve corresponds to an individual mouse.

#### Vaccination elicits partially protective responses against lethal POWV challenge

We next sought to evaluate the protective effects of an ancestral envelope vaccine against a lethal challenge of POWV as a model TBF. Mice were vaccinated with either 10^5^ PFU of SINV (mock-vaccinated) or SINV +E at day 0 and 21 prior to challenge. At 35 days post vaccination, female C57/BL6J mice were challenged with 10^5^ PFU of POWV subcutaneously (SC). Mice that were vaccinated with PBS in place of SINV WT or SINV +E at days 0 and 21 were inoculated with PBS as a mock-challenged control. Mice were monitored over 21 days for survival (**Fig. 5A**), weight (**Fig. 5B**), and clinical score (**Fig. 5C-E**). We observed a significant difference in percent survival in the presence of vaccination against the ancestral envelope antigen with 75% of SINV +E mice surviving to end of study compared to 12.5% survival in mock-vaccinated mice (**Fig. 5A**). Additionally, we observed a recovery of weight after 14 days post infection for vaccinated mice (**Fig. 5B**). We observed that SINV +E vaccinated mice have a delay in disease progression and limited manifestation of disease signs compared to mock-vaccinated mice at day 7 (**Fig. 5D**) and 14 (**Fig. 5E**). This would suggest that SINV +E mice are partially protected against severe disease and mortality during POWV infection.

**Figure 5:**
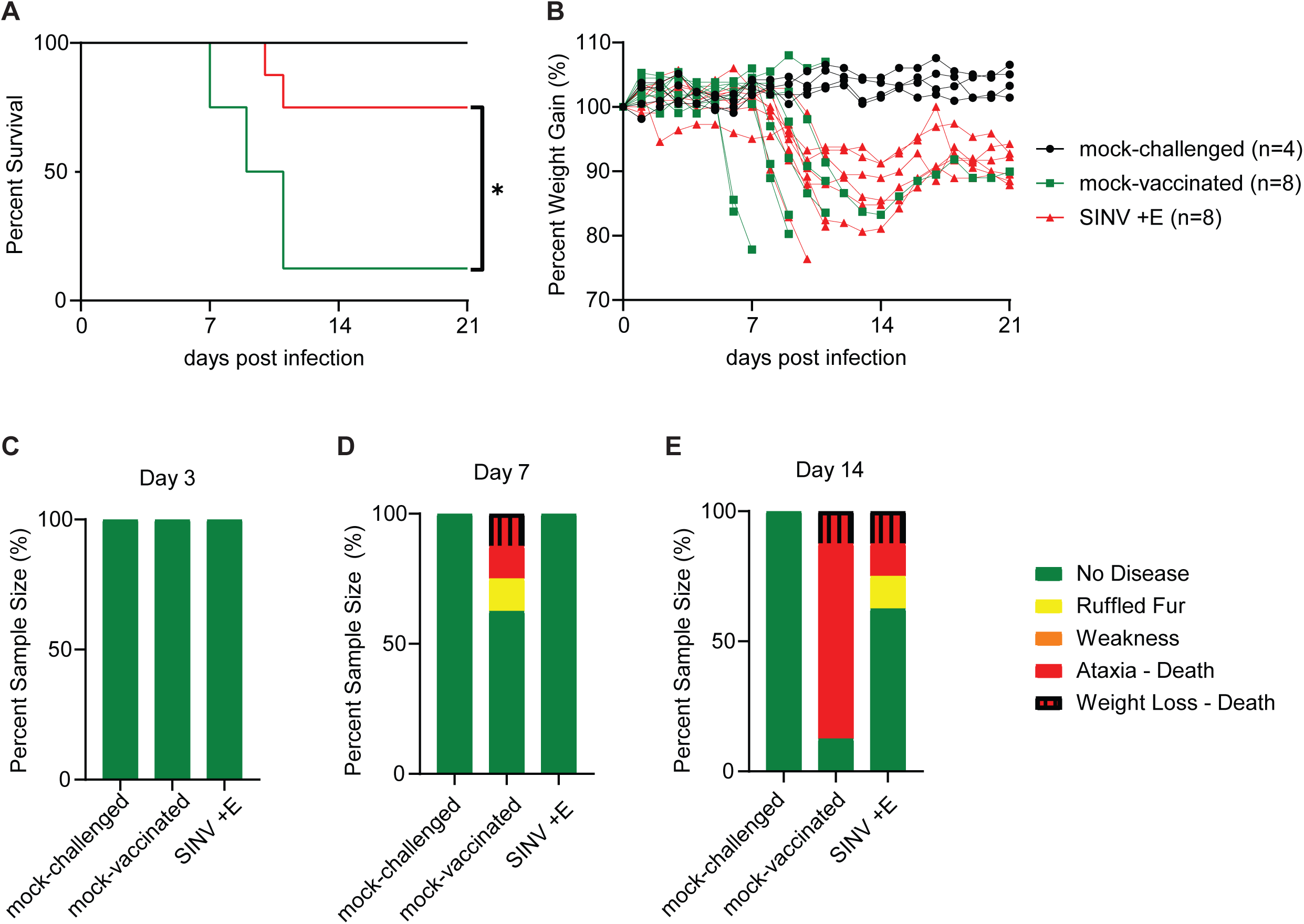
SINV +E vaccination provides partial protection against lethal POWV challenge. (A) Mortality of mock-challenged (black), mock-vaccinated (green), SINV +E vaccinated (red) C57BL/6J mice challenged with 10^5^ PFU of POWV and monitored for 21 days post infection. (B) Weight loss for challenged C57BL/6J mice and monitored for 21 days post infection. (C-E) Clinical scoring results for challenged mice at (C) day 3, (D) day 7, and (E) day 14. Survival of SINV +E vaccinated mice was compared to mock-vaccinated mice with log-rank (Mantel-Cox test, *p<0.05).

### Vaccination against the ancestral envelope antigen provides broader neutralizing capacity but reduced titers compared to a single-target TBF vaccine

The overall goal of this study was to demonstrate the feasibility and advantage of using ASR as a novel approach for antigen design as opposed to current approaches of designing a more specific antigen for a single virus. Previous reports have indicated that there is a low level of cross neutralization for assorted TBFs in individuals seropositive against TBEV [11] or POWV [12]. To address this, we compared our ancestral envelope vaccine to a live-attenuated POWV to assess how broadly protective our SINV +E vaccine candidate compared to a POWV specific vaccine. To generate an attenuated POWV, we introduced a NS4B mutation (M96A) that was shown to attenuate the yellow fever virus 17D vaccine strain [25, 26] into our POWV infectious clone via site-directed mutagenesis and rescued mutant virus. We completed growth curve and plaque morphology analysis on the POWV M96A mutant to assess possible attenuation. Although viral growth kinetics were similar (**Fig. 6A**), POWV-NS4B (M96A) produced plaques that were approximately 61% smaller compared to POWV WT (**Fig. 6B**), suggesting that the NS4B M96A mutation attenuated POWV *in vitro*.

**Figure 6:**
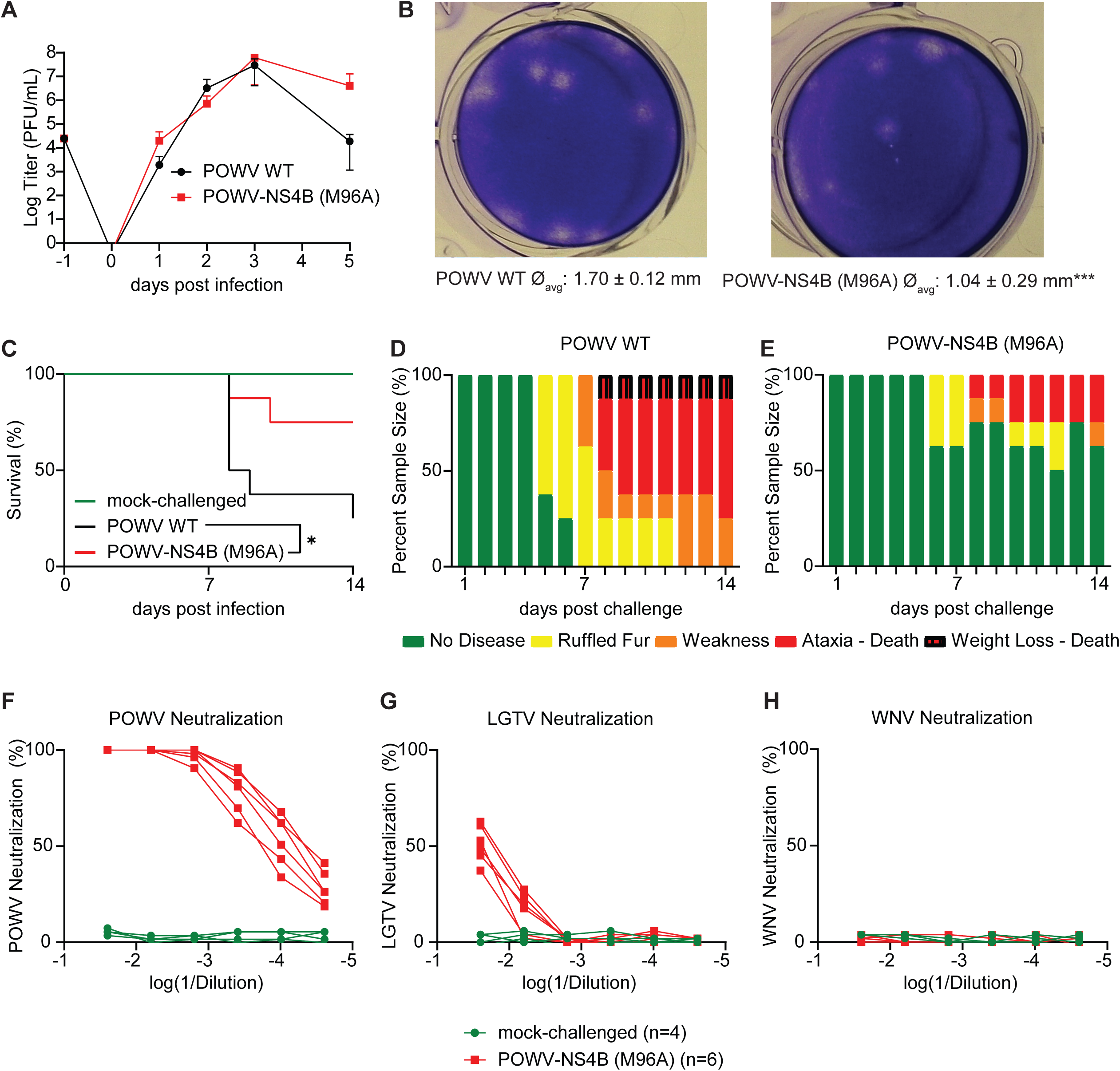
Vaccination against attenuated POWV generates higher neutralizing titers against POWV but lacks broad-spectrum efficacy observed in SINV +E vaccinated mice. (A) Viral titers of passage 1 (p1) from plasmid-transfected BHK-21 cells infected with either virulent (POWV WT) or an attenuated mutant (POWV-NS4B (M96A)). BHK cells were infected at MOI=0.01. Supernatant was collected at indicated times and viral titers were determined by standard plaque assay. (B) Plaque morphology for WT and M96A POWV viruses were obtained at 5-day incubation plaque assays with BHK cells. Average diameter (∅_avg_) is reported for both WT and M96A POWV viruses (***p<0.001, unpaired t-test). (C-H) C57BL/6J mice were challenged with 10^3^ PFU of WT (black) or M96A (red) POWV viruses and monitored over 14 days for survival (C) and clinical score (D-E). Mock-challenged mice (green) were inoculated with PBS as a control. (F-H) Sera was harvested from mock-challenged (black) and POWV-NS4B (M96A)-challenged survivors at 35 days post infection and neutralization was assessed by PRNTs against POWV (F), LGTV (G), and WNV (H).

We next assessed whether the M96A mutation attenuated POWV morbidity and mortality in mice. We challenged female C57BL/6J mice subcutaneously with 10^3^ PFU of either virulent (POWV WT) or attenuated (POWV-NS4B (M96A)) virus diluted in PBS. Mock-challenged mice inoculated with PBS were used as a control. We observed that 25% of POWV WT-challenged mice survived, whereas 75% of POWV-NS4B (M96A) challenged mice survived to end of study (**Fig. 6C**). Additionally, we observed limited disease signs for POWV-NS4B (M96A) challenged mice compared to the POWV WT challenged group over 14 days (**Fig. 6D-E**), suggesting that POWV-NS4B (M96A) is attenuated *in vivo*.

We then assessed the breadth of neutralization of sera harvested from POWV-NS4B (M96A) challenged mice. We assessed neutralizing titers of sera against POWV (**Fig. 7F**), LGTV (**Fig. 7G**), and WNV (**Fig. 7H**) at day 35 post infection. Interestingly, we observed that while the neutralizing titers against POWV are higher than our ancestral envelope vaccinated mice, the broad-spectrum efficacy was lost as there was reduced neutralization observed against LGTV.

## DISCUSSION

In this project we tested a novel and effective approach for designing a broad-spectrum vaccine against tick-borne flaviviruses. We demonstrated that a vaccine candidate based on ancestral sequence reconstruction (ASR) is safe with no adverse effects in mice vaccinated using a live alphavirus platform. We showed that our vaccine platform expressing a common ancestral envelope antigen (envelope (E)) induces antibodies that can effectively neutralize at least two phylogenetically distinct TBFs (POWV and LGTV) at comparable titers, whereas vaccination using an attenuated POWV generated strong POWV-specific responses with limited cross-reactivity. Furthermore, we demonstrated that vaccination against our reconstructed ancestral envelope antigen significantly reduces mortality and disease in a mouse model. Taken together, our findings indicate that our ASR approach is not only viable but effective in protecting against TBF viral challenge.

Like any vaccine development approach, additional factors must be considered to optimize the level of protection necessary for effective protection. In the current study we assessed vaccine efficacy within a standard dose schedule; however, it is rational to assume that adjustments in administration and dosage may influence efficacy reported. The recommended vaccine available in the U.S.A. against TBE (TicoVac®) is an inactivated formula that requires 3 doses over the span of 12-18 months [27], with additional doses required for individuals over 50 years of age. Alterations in vaccine dosage and administration schedule may influence overall immunogenicity and level of protection achievable by our ASR-generated antigen. Additionally, the platform used for antigen expression may also influence vaccine efficacy. For this study, we utilized a well-established alphavirus-based system for antigen expression. However, as noted in previous studies, there are limitations to this approach long term. These concerns include overall safety in using a self-replicating viral platform [28] and stability of inserted genes following subgenomic promoters [23]. While we can conclude that no adverse effects were observed in mice vaccinated at doses upwards to 10^5^ PFU, stability of antigen expression long term may need to be considered. Alternative expression systems such as an mRNA platform that has been previously used successfully against POWV [12] may reduce potential stability issues.

Although the results of this study are promising and account for antigenic diversity within the TBF clade to generate broad immunological responses, the ASR method is reliant on sampled virus sequences selected as representative strains for their viral species (listed in Table S1 for this study). As a result, the ASR approach may introduce unintentional sequence bias that can affect the reconstruction of ancestral sequences and may result in the production of a suboptimal antigen sequences. Inclusion of more representative strains and species may overcome these hinderances and improve overall efficiency in neutralizing titers.

Further study is necessary to clarify the efficacy of an ancestral antigen-based approach against other TBFs that were beyond our ability to assess. Specifically, additional information is needed to determine how effective our vaccine is against the more virulent TBFs that require BSL4 facilities. Based on our assessment of neutralizing titers against a LGTV-TBEV chimera (**Fig. 4C**), we can assume that this current vaccine construct would have limited, if any, neutralizing titers against wildtype TBEV. Additionally, we observed lower neutralizing titers in SINV +E vaccinated mice against POWV but higher against LGTV compared to our attenuated POWV vaccine characterized in Figure 6. This observation leads us to conclude that while our ASR-derived vaccine has greater sensitivity against multiple TBFs, its specificity may be reduced in comparison. A possible approach to address this includes evaluating the immunogenicity of antigens of internal nodes closer to the modern-day clades of interest. For example, a vaccine expressing the antigen derived from node 20 in the TBF phylogenetic tree (**Fig. 1A**) may improve neutralizing titers against the more virulent TBFs. In summary, we provide the first demonstration that an ASR-derived antigen to a TBF envelope protein provides effective immunity against multiple modern-day TBFs. Our findings provide foundational understanding regarding the potential of ASR as a novel antigen design tool that can be used in vaccine design and potentially alleviate disease burden to those most at risk.

## MATERIALS AND METHODS

### Phylogenetic analysis and generation of ancestral sequence

Ancestral sequences were calculated from amino acid alignments used to generate an E phylogenetic tree using MEGA11 (version 11.0.3) [21]. Sequences from multiple modern-day tick-borne flaviviruses were obtained from GenBank and phylogenetic relationship was inferred by using the Maximum Likelihood (ML) method and a Jones-Taylor-Thornton (JTT) matrix-based model. AlphaFold2 was used to model and visualize the protein structure of the predicted ancestral TBF E amino acid sequence derived from ASR [22]. Sequences used for phylogenetic analysis and ancestral sequence reconstruction are provided in Table S1.

### Viruses, cells, and animal models

POWV-Spooner (lineage II) infectious clone was provided by A. Brault (Centers for Disease Control and Prevention, USA) and rescued and passaged once on BHK-21 cells (Syrian golden hamster kidney fibroblasts) (ATCC CCL-10) as previously described [29]. Rescued LGTV-TP21 and LGTV-TBEV chimeric infectious clones were provided by A. Överby (Umeå University, Sweden) and passaged once on VeroB4 cells (African green monkey kidney epithelial cells) or A549 *Mavs-/-* cells, respectively, as described [24, 30]. WNV-FtC-3699 was passaged once on Vero CCL-81 cells (ATCC CCL-81). Supernatants of these cultures were aliquoted and frozen at -80°C. Infectious virus titer was measured using a standard plaque assay. BHK-21 and Vero cells were maintained in DMEM supplemented with 7% fetal bovine serum (FBS) and 1% penicillin/streptomycin at 37 °C and 5% CO_2_.

Female wildtype C57BL/6J (B6, 000664) mice were purchased from Jackson Laboratories (Bar Harbor, Maine, USA) and housed in Colorado State University ABSL-3 containment, according to the animal protection act. Approval for animal protocols was obtained by the Colorado State University Institutional Animal Care and Use Committee (protocol #3394; #7224).

### Plasmid construction

The alphavirus plasmid construct pBG167 (SINV WT) was generated and characterized as described [23]. An ancestral TBF E (ASR TBF E) antigen gBlock was designed and purchased through Integrated DNA Technologies (IDT). Codon biases were considered and optimized using the IDT Codon Optimization Tool. The gBlock was inserted using the NEBuilder^®^ HiFi DNA Assembly Cloning Kit (New England Biolabs, E5520S) according to manufacturer instructions. Briefly, SINV WT plasmid was digested with *XbaI* restriction enzyme (RE) (New England Biolabs, R0145S) to generate a linear product and gel extracted. The gBlock underwent PCR amplification and extension using Q5^®^ High-Fidelity DNA Polymerase (New England Biolabs, M0491S) (forward: ACAACACCACCACCTATGTTTTTTGCAGTGACGGCTCT; reverse: CGTCTAGGATCCATGGTTTACGCCCCAACTCCCCT). NEBuilder HiFi DNA Assembly (New England Biolabs) was completed in a single reaction consisting of both SINV WT RE digested vector and PCR-amplified gBlock insert (DNA molar ratio of 1:2) and 10 μL NEBuilder HiFi DNA Assembly Master Mix for a total reaction of 20 μL. Samples were mixed via pipetting and incubated at 37°C for 1 hour, then the reactions were transformed into XL-10 *E. coli* cells and selected on LB agar plates with 50ug/ml ampicillin. Isolated colonies were selected and plasmid DNA was purified using a QIAprep Spin Miniprep Kit (Qiagen, 27106) according to manufacturer instructions. Plasmid sequences were confirmed via whole plasmid sequencing (Plasmidsaurus, Louisville, Kentucky, USA).

### Rescue and characterization of infectious clones

BHK-21 cells were plated the day prior to transfection into 6 well plates at approximately 60-70% confluency. Cells were transfected with 125 ng plasmid DNA using Lipofectamine 3000 (Invitrogen, L3000001) following manufacturer recommendations. Transfection media was removed and replaced with fresh media 4 hours post-transfection. Supernatant containing virus was harvested at 4 days post transfection when >50% CPE was observed. This initial passage stock (p0) was aliquoted and stored at -80°C. Viral stocks were passaged once in BHK-21 cells (p1), titered by plaque assay, and stored at -80°C in single-use aliquots to serve as vaccine stocks for the study.

### Plaque assays

Standard plaque assays were used to quantify infectious virus. Briefly, either BHK-21 (SINV, POWV, LGTV, LGTV-TBEV) or Vero (WNV) cells were plated the day prior to infection at 90-100% confluency. Virus was serially diluted, added onto cell monolayers, and incubated at 37°C for 1 hour. Cells were overlaid with a semisolid tragacanth medium and incubated for 3 (SINV, WNV), 4 (LGTV, LGTV-TBEV) or 5 (POWV) days. Overlay medium was removed, cells were fixed with 20% ethanol and subsequently stained with 0.1% crystal violet. Plaques were counted manually.

### Growth curves

BHK-21 cells were plated the day prior to infection at 2.5x10^5^ cells/mL in 6 well plates. Cells were infected with virus (MOI=0.01) diluted in infection media (DMEM supplemented with 2% FBS and 1% penicillin/streptomycin) for 1 hour at 37°C. Cells were washed 3 times with PBS and fresh growth media was added. Supernatant was sampled daily with removed volume replaced with fresh media, and viral titers determined with plaque assays.

### qRT-PCR

RNA was extracted using the QIAamp Viral RNA Mini Kit (Qiagen, 52904) according to manufacturer’s instructions. qRT-PCR was performed using EXPRESS One-Step SYBR GreenER kit (Invitrogen, 11780200) according to the manufacturer’s instructions with primers targeting the ancestral TBF E insert (forward: CGAGACCCGAGAGTATTGCC; reverse: ACCGTGAACGATGCTGTCTT).

### Vaccination and POWV challenge

Nine-week old C57BL/6J mice were vaccinated subcutaneously (SC) in the right quadricep with 10^5^ PFU of either control (SINV WT) or envelope antigen expressing (SINV +E) vaccine stocks diluted into 50 μL of PBS. Mock-vaccinated mice were inoculated with 50 μL PBS SC. Twenty-one days later mice were boosted with an identical formulation.

For evaluation of the ancestral TBF E vaccine studies, mice were inoculated SC with a lethal challenge of 10^5^ PFU of POWV diluted in 50 μL of PBS. For comparison against an attenuated POWV construct, mice were infected SC with 10^3^ PFU of either POWV WT or POWV-NS4B mutant diluted in 50 μL of PBS. Control mock challenged mice were inoculated SC with 50 μL PBS instead of POWV. Mice were monitored over 35 days to assess vaccine safety prior to viral challenge. Following lethal challenge, mice were monitored 14-21 days post-challenge for weight loss, signs of neurological disease, and mortality. The clinical scoring system used was as follows: 0=healthy, 1=ruffled fur, hunched back, 2=hindlimb weakness, 3=ataxia, tremors, 4=paralysis, 5=deceased. Mice that reached a clinical score of 3 or exhibited weight loss greater than 20% compared to starting weight were humanely euthanized.

### Sera harvesting and isolation

Sera was harvested prior at 35 days post initial vaccination prior to viral challenge via cardiac puncture. Briefly, mice were anesthetized using isoflurane vapor (Kent Scientific Corporation, SomnoFlo, SF-01) until sedated. Approximately 400 to 600 μL of whole blood was collected in serum separation tubes (BD Microtainer® #365967). Collection tubes were inverted and incubated at room temperature for 30 minutes for a blood clot to form. Samples were centrifuged at 1300 x g for 10 minutes at 4 °C. Sera was then separated into a new 1.5 mL Eppendorf tube, heat inactivated at 56 °C for 30 minutes, aliquoted, and stored at -80°C for later analysis.

### Plaque reduction neutralization tests (PRNTs)

A standard PRNT was performed on POWV, LGTV, LGTV-TBEV, and WNV to assess antibody neutralization titers as described [31]. Briefly, either BHK-21 (POWV, LGTV, LGTV-TBEV) or Vero cells (WNV) were plated 1 day prior to infection. A serial dilution of heat-inactivated sera was incubated with approximately 50 PFU of virus for 1 hour at 37°C. Following incubation, the serum:virus inoculum was added to cells, incubated 1 hour at 37°C, then overlaid with semisolid tragacanth medium. Plates were incubated at 37°C and 5% CO_2_ for 3 (WNV), 4 (LGTV, LGTV-TBEV), or 5 (POWV) days. Cells were fixed and stained with 20% ethanol and 0.1% crystal violet and plaques were quantified manually. PRNT_50_ values were calculated in GraphPad Prism 10.

### Generation and characterization of attenuated POWV-NS4B (M96A)

Attenuation of the virulent POWV-Spooner infectious clone (POWV WT) was achieved by identifying conserved residues shared between POWV WT and virulent yellow fever virus (GenBank: NP_041726.1) by multiple sequence alignment. Selection of candidate residues for mutagenesis was based on known residues associated with attenuation within the vaccine strain yellow fever 17D. A single point mutation (NS4B M96A) was introduced into the POWV infectious clone plasmid using the Q5^®^ Site-Directed Mutagenesis Kit (New England Biolabs, E0554S) according to manufacturer recommendations (forward: ACACGTTGTGgcaCTTGGCGTGACTTC; reverse: CCAGCCACGCCAAAGAAT). The resulting infectious clone was rescued and characterized as previously mentioned. Plaque morphology and size was evaluated for 8-10 plaques fixed and stained after 5 days of incubation using ImageJ and statistically evaluated using GraphPad Prism 10.

### Statistical analysis

All statistical analyses were performed using GraphPad Prism 10 (version 10.5.0). Statistical differences in survival curves were determined using a Mantel-Cox test. Statistical significance for qRT-PCR by student’s unpaired t-test. Data for growth curves in tissue culture is representative of three independent experiments and three biological replicates. Data for survival and PRNT are representative of two independent experiments (n=4-8). Statistical significance has been indicated within the figures with asterisks (*p < 0.05, **p < 0.01, ***p < 0.001).

## DATA AVAILABILTIY

All data reported in this paper will be shared by the lead contact upon request. This paper does not report original code. Any additional information required to reanalyze the data reported in this paper is available from the lead contact upon request.

## ACKNOWLEDGEMENTS

This research was supported by the U.S. Department of Defense (DoD) (award no. W81XWH-22-1-0905). The funder had no role in study design, data collection and analysis, decision to publish, or preparation of the manuscript.

## AUTHOR CONTRIBUTIONS

Experimental conceptualization, C.E.T., B.J.G., G.D.E.; data curation, C.E.T., B.J.G., G.D.E.; formal analysis, C.E.T., B.M.E., M.J.S., B.J.G., G.D.E.; funding acquisition, B.J.G., G.D.E.; investigation, C.E.T., B.M.E., M.J.S., B.J.G., G.D.E.; methodology, C.E.T., B.J.G., G.D.E.; project administration, C.E.T.; resources, B.J.G., G.D.E.; software, C.E.T.; supervision, C.E.T., B.J.G., G.D.E.; validation, C.E.T., B.M.E., M.J.S., B.J.G., G.D.E.; visualization, C.E.T., B.J.G., G.D.E.; writing – original draft, C.E.T.; writing – review & editing, C.E.T., B.M.E., M.J.S., B.J.G., G.D.E., who reviewed and approved the content and submission of the final manuscript.

---

**Fig. S1:**
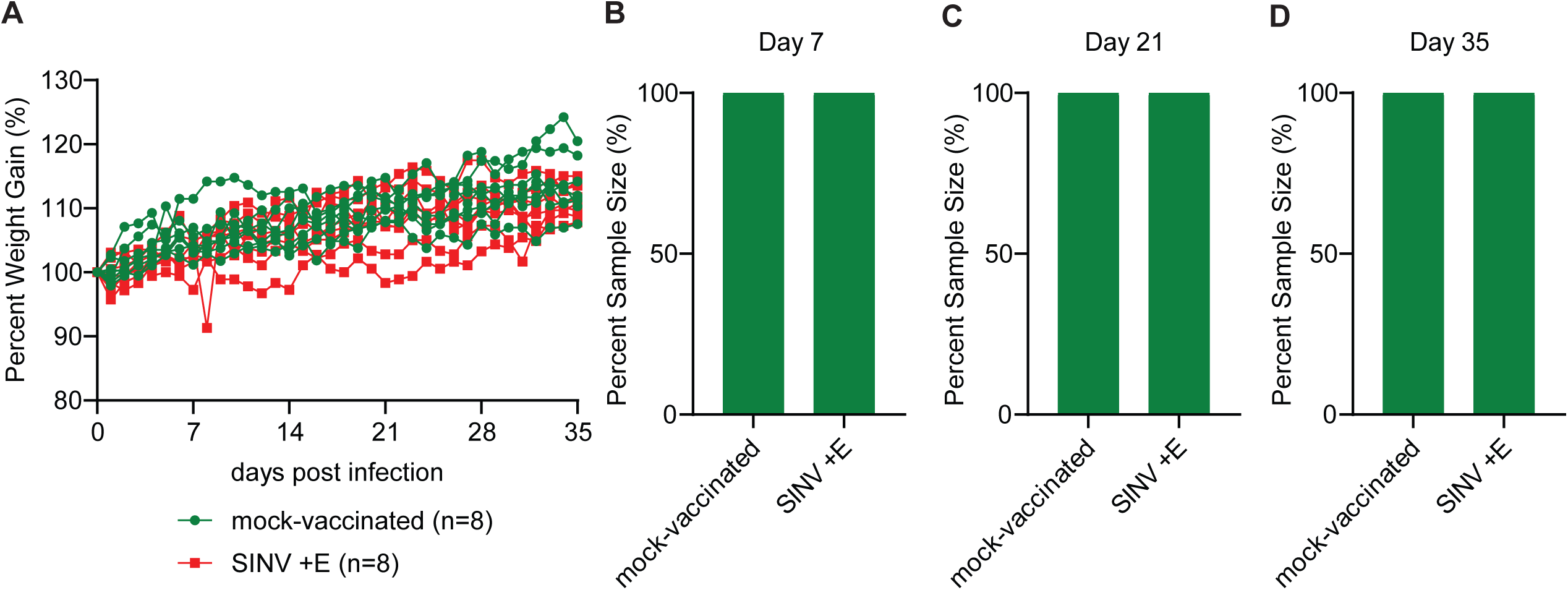
Weight (A) and (B-D) clinical signs of SINV +E vaccinated C57/BL6J mice at day 7 (B), 21 (C), and 35 (D) post inoculation.

**Table S1:**
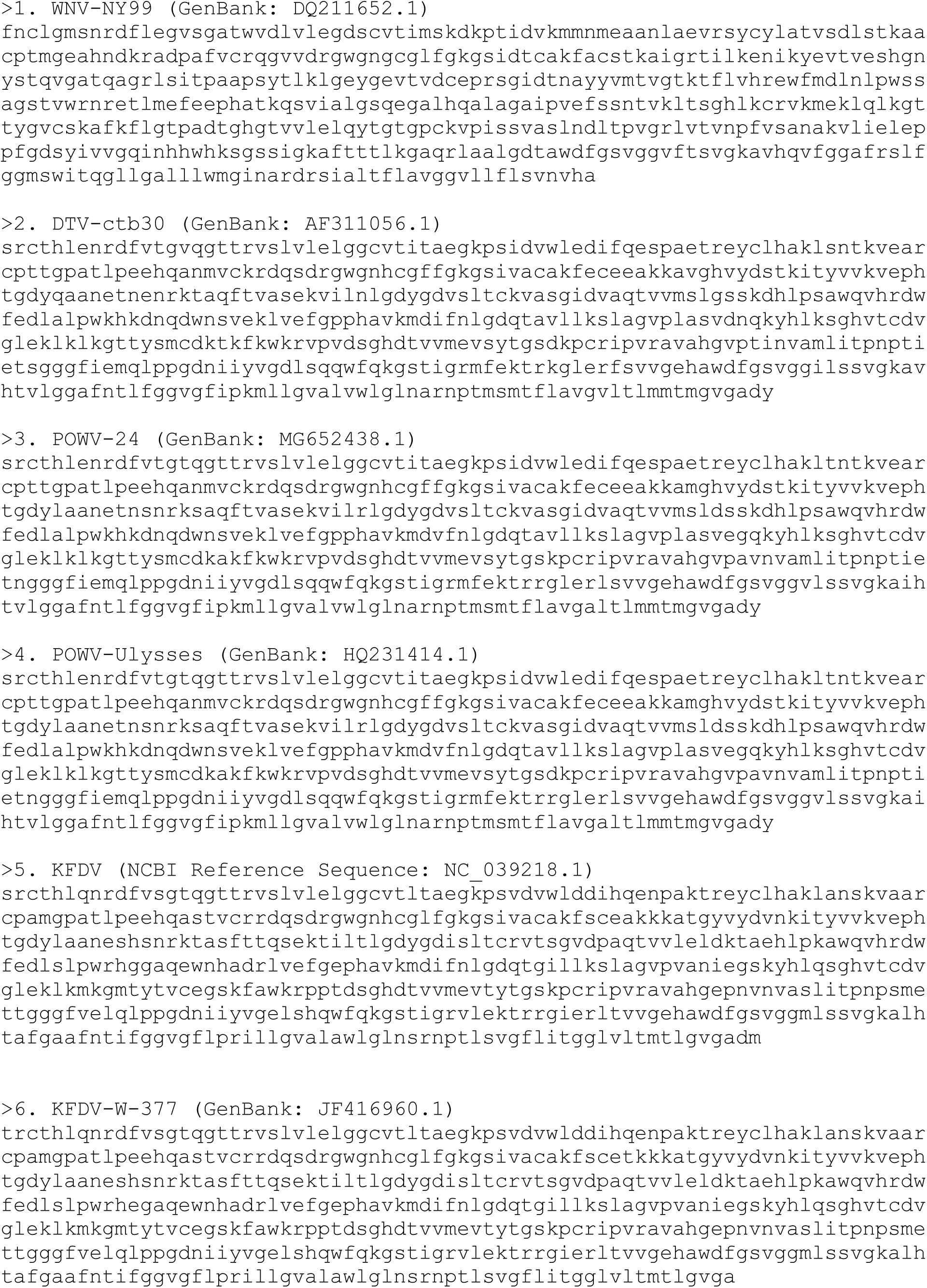
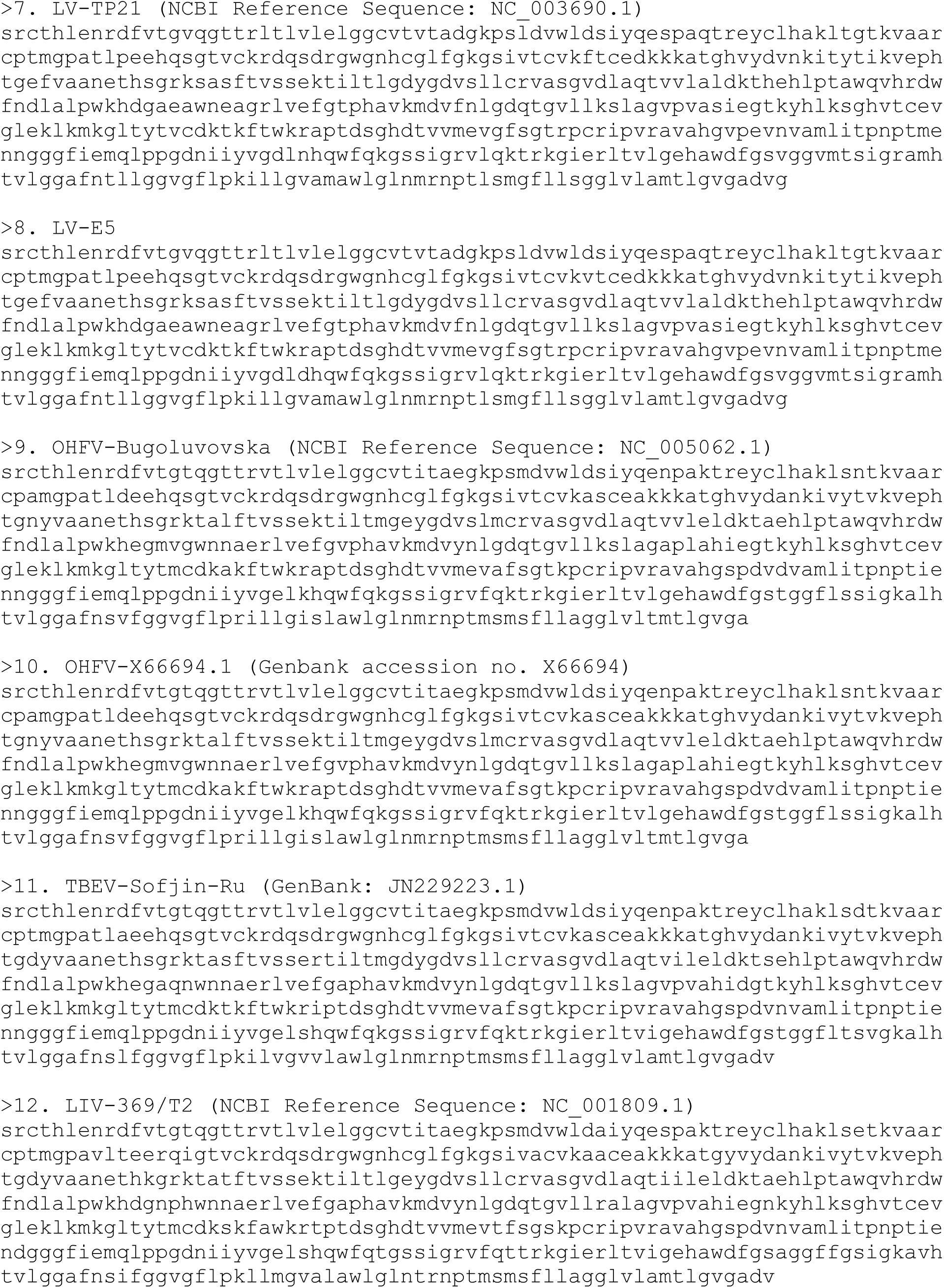
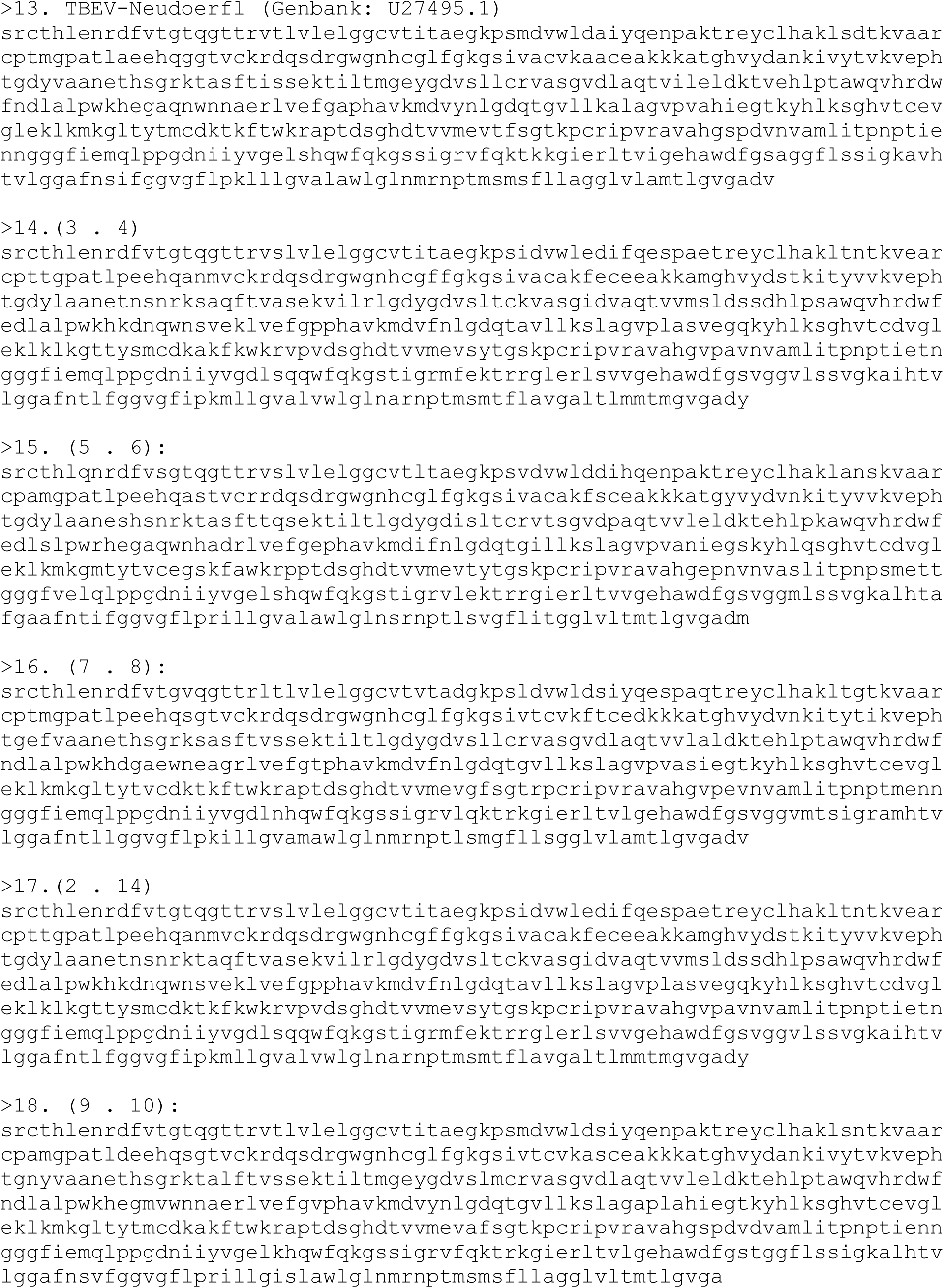
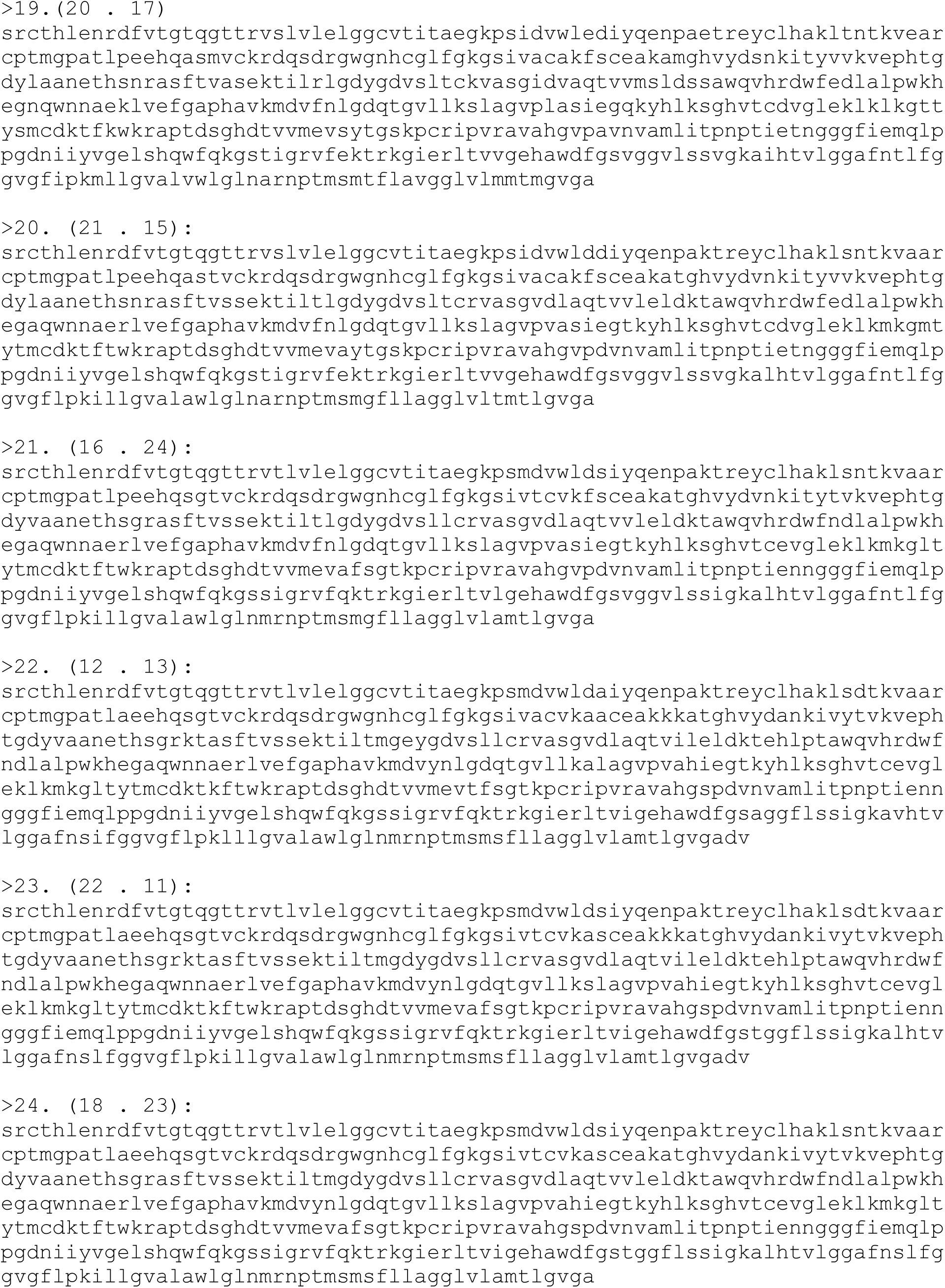
Virus sequences used and reconstructed for ancestral sequence reconstruction and antigen design for Figure 1.

## REFERENCES

1. CDC. Data and Maps for Powassan. Powassan Virus. 2025. https://www.cdc.gov/powassan/data-maps/index.html. Accessed 6 Oct 2025.

2. McCluskey JM, Wichman CS, Degarege A, Brett-Major D. Leveraging Lyme Disease Surveillance to Simulate Powassan Virus Prevalence. J Infect Dis. 2025;:jiaf368. 10.1093/infdis/jiaf368.

3. Siegel E, Xu G, Killinger P, Brown CM, Rich SM. Passive surveillance of Powassan virus in human-biting ticks and health outcomes of associated bite victims. Clin Microbiol Infect. 2024;30:1332–4. 10.1016/j.cmi.2024.06.012.

4. Kwasnik M, Rola J, Rozek W. Tick-Borne Encephalitis-Review of the Current Status. J Clin Med. 2023;12:6603. 10.3390/jcm12206603.

5. Telford SR III, Stewart PE, Bloom ME. Increasing Risk for Tick-Borne Disease: What Should Clinicians Know? JAMA Intern Med. 2024;184:973–4. 10.1001/jamainternmed.2024.1754.

6. Damian D. The Growing Threat of Tick-Borne Viruses: Global Trends, Clinical Outcomes, and Diagnostic Strategies. Viral Immunol. 2025;38:125–36. 10.1089/vim.2025.0019.

7. Petit MJ, Johnson N, Mansfield KL. Vectorial dynamics underpinning current and future tick-borne virus emergence in Europe. J Gen Virol. 2024;105:002041. 10.1099/jgv.0.002041.

8. Bouchard C, Dibernardo A, Koffi J, Wood H, Leighton P, Lindsay L. N Increased risk of tick-borne diseases with climate and environmental changes. Can Commun Dis Rep. 2019;45:83–9. 10.14745/ccdr.v45i04a02.

9. Kubinski M, Beicht J, Gerlach T, Volz A, Sutter G, Rimmelzwaan GF. Tick-Borne Encephalitis Virus: A Quest for Better Vaccines against a Virus on the Rise. Vaccines. 2020;8:451. 10.3390/vaccines8030451.

10. Hansson KE, Rosdahl A, Insulander M, Vene S, Lindquist L, Gredmark-Russ S, et al. Tick-borne Encephalitis Vaccine Failures: A 10-year Retrospective Study Supporting the Rationale for Adding an Extra Priming Dose in Individuals Starting at Age 50 Years. Clin Infect Dis Off Publ Infect Dis Soc Am. 2020;70:245–51. 10.1093/cid/ciz176.

11. McAuley AJ, Sawatsky B, Ksiazek T, Torres M, Korva M, Lotrič-Furlan S, et al. Cross-neutralisation of viruses of the tick-borne encephalitis complex following tick-borne encephalitis vaccination and/or infection. NPJ Vaccines. 2017;2:5. 10.1038/s41541-017-0009-5.

12. VanBlargan LA, Himansu S, Foreman BM, Ebel GD, Pierson TC, Diamond MS. An mRNA Vaccine Protects Mice against Multiple Tick-Transmitted Flavivirus Infections. Cell Rep. 2018;25:3382–3392.e3. 10.1016/j.celrep.2018.11.082.

13. Liberles DA. Ancestral Sequence Reconstruction. OUP Oxford; 2007.

14. Straub K, Merkl R. Ancestral Sequence Reconstruction as a Tool for the Elucidation of a Stepwise Evolutionary Adaptation. In: Sikosek T, editor. Computational Methods in Protein Evolution. New York, NY: Springer; 2019. p. 171–82. 10.1007/978-1-4939-8736-8_9.

15. Selberg AGA, Gaucher EA, Liberles DA. Ancestral Sequence Reconstruction: From Chemical Paleogenetics to Maximum Likelihood Algorithms and Beyond. J Mol Evol. 2021;89:157–64. 10.1007/s00239-021-09993-1.

16. Furukawa R, Toma W, Yamazaki K, Akanuma S. Ancestral sequence reconstruction produces thermally stable enzymes with mesophilic enzyme-like catalytic properties. Sci Rep. 2020;10:15493. 10.1038/s41598-020-72418-4.

17. Prakinee K, Phaisan S, Kongjaroon S, Chaiyen P. Ancestral Sequence Reconstruction for Designing Biocatalysts and Investigating their Functional Mechanisms. JACS Au. 2024;4:4571–91. 10.1021/jacsau.4c00653.

18. Ducatez MF, Bahl J, Griffin Y, Stigger-Rosser E, Franks J, Barman S, et al. Feasibility of reconstructed ancestral H5N1 influenza viruses for cross-clade protective vaccine development. Proc Natl Acad Sci U S A. 2011;108:349–54. 10.1073/pnas.1012457108.

19. Ross HA, Nickle DC, Liu Y, Heath L, Jensen MA, Rodrigo AG, et al. Sources of variation in ancestral sequence reconstruction for HIV-1 envelope genes. Evol Bioinforma Online. 2007;2:53–76.

20. Gaschen B, Taylor J, Yusim K, Foley B, Gao F, Lang D, et al. Diversity Considerations in HIV-1 Vaccine Selection. Science. 2002;296:2354–60. 10.1126/science.1070441.

21. Tamura K, Stecher G, Kumar S. MEGA11: Molecular Evolutionary Genetics Analysis Version 11. Mol Biol Evol. 2021;38:3022–7. 10.1093/molbev/msab120.

22. Jumper J, Evans R, Pritzel A, Green T, Figurnov M, Ronneberger O, et al. Highly accurate protein structure prediction with AlphaFold. Nature. 2021;596:583–9. 10.1038/s41586-021-03819-2.

23. Steel JJ, Henderson BR, Lama SB, Olson KE, Geiss BJ. Infectious alphavirus production from a simple plasmid transfection. Virol J. 2011;8:356. 10.1186/1743-422X-8-356.

24. Rosendal E, Bisikalo K, Willekens SMA, Lindgren M, Holoubek J, Svoboda P, et al. Influence of the pre-membrane and envelope proteins on structure, pathogenicity, and tropism of tick-borne encephalitis virus. J Virol. 2025;99:e00870–25. 10.1128/jvi.00870-25.

25. Davis EH, Beck AS, Strother AE, Thompson JK, Widen SG, Higgs S, et al. Attenuation of Live-Attenuated Yellow Fever 17D Vaccine Virus Is Localized to a High-Fidelity Replication Complex. mBio. 2019;10:10.1128/mbio.02294-19. 10.1128/mbio.02294-19.

26. Beck AS, Wood TG, Widen SG, Thompson JK, Barrett ADT. Analysis By Deep Sequencing of Discontinued Neurotropic Yellow Fever Vaccine Strains. Sci Rep. 2018;8:13408. 10.1038/s41598-018-31085-2.

27. CDC. Tick-borne Encephalitis Vaccine Information for Healthcare Providers. Tick-borne Encephalitis Virus. 2024. https://www.cdc.gov/tick-borne-encephalitis/hcp/vaccine/index.html. Accessed 8 Oct 2025.

28. Maine CJ, Miyake-Stoner SJ, Spasova DS, Picarda G, Chou AC, Brand ED, et al. Safety and immunogenicity of an optimized self-replicating RNA platform for low dose or single dose vaccine applications: a randomized, open label Phase I study in healthy volunteers. Nat Commun. 2025;16:456. 10.1038/s41467-025-55843-9.

29. Kenney JL, Anishchenko M, Hermance M, Romo H, Chen C-I, Thangamani S, et al. Generation of a Lineage II Powassan Virus (Deer Tick Virus) cDNA Clone: Assessment of Flaviviral Genetic Determinants of Tick and Mosquito Vector Competence. Vector-Borne Zoonotic Dis. 2018;18:371–81. 10.1089/vbz.2017.2224.

30. Chotiwan N, Rosendal E, Willekens SMA, Schexnaydre E, Nilsson E, Lindqvist R, et al. Type I interferon shapes brain distribution and tropism of tick-borne flavivirus. Nat Commun. 2023;14:2007. 10.1038/s41467-023-37698-0.

31. Gallichotte EN, Fitzmeyer EA, Williams L, Spangler MC, Bosco-Lauth AM, Ebel GD. WNV and SLEV coinfection in avian and mosquito hosts: impact on viremia, antibody responses, and vector competence. J Virol. 98:e01041–24. 10.1128/jvi.01041-24.

